# *Hortaea werneckii* isolates exhibit different pathogenic potential in the invertebrate infection model *Galleria mellonella*

**DOI:** 10.1101/2022.10.24.513617

**Authors:** Stephanie Anthonies, José M. Vargas-Muñiz

## Abstract

*Hortaea werneckii* is a black yeast with a remarkable tolerance to salt. Most studies have been dedicated to understanding how *H. werneckii* adapt to hypersaline environments. *H. werneckii* has an unconventional cell cycle in which it alternates between fission and budding, which is modulated by cell density. Additionally, *H. werneckii* can cause superficial mycosis of the palm and sole of humans. Here, we determine the impact of salt concentration on the EXF-2000 strain’s cell division pattern and morphology. At low density and no salt, EXF-2000 primarily grows as pseudohyphae dividing mainly by septation. When grown in the presence of salt at a similar concentration to saltwater or hypersaline environments, we observe it grows by first by undergoing fission followed by budding at the poles. Then, we examined a collection of 16 isolates, including isolates from marine and hypersaline environments and isolates from patients. Isolates exhibit a wide diversity in colony shape and cellular morphology. The isolates grew as yeast, pseudohyphae, and true hyphae, indicating that isolates can exhibit various cell morphologies under similar environmental conditions. We used the insect larvae *Galleria mellonella* to determine the pathogenic potential of our isolates. We observe that only a subset of isolates can cause death in our model, and there was no correlation between *H. werneckii* morphology and capacity to cause disease. Taken together, *H. werneckii* genomic and phenotypic diversity can serve as a model to better understand how phenotypes and pathogenic potential evolve in environmental fungi.

## Introduction

Morphogenetic transition is a crucial feature of fungal pathogenesis (Zaragoza and Nielsen, 2013; Kornitzer, 2019; Gupta et al., 2021). Fungal pathogens alter their morphology in reaction to their host environment, facilitating host-tissue invasion and disease progression (Zaragoza and Nielsen, 2013; Kornitzer, 2019). In the case of the human pathogen *Candida albicans*, the yeast hyphal transition is indispensable for pathogenesis (Kornitzer, 2019). In *C. albicans*, strains locked in either the hyphal or yeast morphology have attenuated virulence compared to the wild-type strain. When haploid yeasts of the basidiomycete *Cryptococcus neoformans* are in the lungs of the human host, they transition into a large, polyploid cell known as Titan cells (Zaragoza and Nielsen, 2013; Dambuza et al., 2018; Hommel et al., 2018). Titan cells account for 20% of the *C. neoformans* cells in the lungs and influence the fungal interactions with its host (Okagaki et al., 2010). They can lead to persistence in the host and increased drug resistance. The human fungal pathogens *Histoplasma capsulatum, Blastomyces dermatiditis, Coccidioides immitis, Coccidioides posadassi, Talaromyces mernefeii, Paracoccidioides brasiliensis, Paracoccidiodes lutzii*,and *Sporothrix schenckii*have thermally regulated dimorphism. These fungi grow predominantly as hyphae in the environment. However, inside the host, these organisms undergo a host-adapted developmental cycle that leads to the production of yeast or spherules (*B. dermatiditis*). Interestingly, most of these fungi are primarily human pathogens that cause endemic infections in healthy humans. Plant fungal pathogens also depend on morphogenetic transition for full virulence. When the rice blast fungus *Magnaporthe oryzae* spores land on a leaf, they break dormancy and form a germ tube. The germ tube then differentiates into a dome structure known as the appressorium(Wang et al., 2005; Gupta et al., 2021). The appressorium provides the turgor pressure that allows the penetration peg to puncture the leaf cuticles; mutants defective on appressoria formation have attenuated virulence(Saunders et al., 2010; Dagdas et al., 2012; Kunova et al., 2013). Thus, being able to alter their morphology in response to their host is essential for fungal pathogenesis.

*Hortaea werneckii (Ascomycota, Capnodiales*) is an extremely halotolerant yeast that can withstand up to 5M NaCl (Plemenitaš et al., 2014). *H. werneckii* has been isolated from salterns, seawater, coral, sponges, deep-sea sediments, and the Atacama Desert (Chile) (Gostinčar et al., 2018). Most of *H. werneckii* studies have focused on their extraordinary ability to survive at high salt concentrations (Petrovič et al., 2002; Turk et al., 2004; Kogej et al., 2007; Lenassi et al., 2007; Vaupotix and Plemenitaš, 2007). Nonetheless, *H. werneckii* can cause a superficial infection of the human hands and feet known as tinea nigra (Ng et al., 2005; Bonifaz et al., 2008; Giordano Lorca et al., 2018). Tinea nigra infections frequently occur in countries with tropical climates, and in rare cases can cause systemic infection known as disseminated phaeohyphomycosis (Bonifaz et al., 2013).

Most *H. werneckii* isolates are results of intraspecific hybridization of two haploid *H. werneckii*,which may contribute to phenotypical diversity (Gostinčar et al., 2018; Zalar et al., 2019; Romeo et al., 2020). Remarkably, *H. werneckii* cells can grow via both fission and budding; their cells grow from both poles (similar to *Schizosaccharomyces pombe*) and are divided by a median septum (Mitchison-Field et al., 2019). After the septation event occurs, cells start budding from both poles. Daughter cells then follow this same pattern of septation followed by budding. Cell size and division time were highly heterogeneous when compared to those of well-studied yeasts *Saccharomyces cerevisiae* and *S. pombe. H. wereneckii* cell division and morphology is dependent on cell density (Goshima, 2022).

Here, we describe how the environment influences *H. werneckii* cell division and morphology. We characterized the growth pattern of *H. werneckii* EXF-2000 isolate, which has a sequenced genome. We observed that the concentration of sodium chloride dictates whether EXF-2000 grows predominantly with a pseudohyphal-like or yeast morphology. We also observed that septation occurs more often in low NaCl concentrations. Due to this morphogenetic plasticity, we characterized the macroscopic and microscopic morphology of different *H. werneckii* isolates.

When grown in the presence of marine salts, *H. werneckii* exhibited a variety of morphology (from yeast to hyphae). We also observed different levels of tolerance to the antifungal drugs fluconazole and caspofungin. To correlate if morphology influences pathogenic potential, we established a *Galleria mellonella* pathogenesis model for *H. werneckii*. We observed that pathogenic potential varies between strains but was not correlated to morphological capacity. Based on these results, we suggest that *H. werneckii* intraspecific hybridization generates phenotypic diversity, thus, allowing this species to adapt to a myriad of environments, including the human body.

## Materials and Methods

### Strains, Media, Culture conditions

Previously isolated *Hortaea werneckii* and *Salinomyce thailandica* were used for these studies (Table 1). Unless otherwise specified, yeast cells were grown in YPD supplemented with Instant Ocean (Instant Ocean YPD) [yeast extract 10 g/L, peptone 20 g/L, dextrose 20 g/L, Instant Ocean (Spectrum Brands, Blacksburg, VA) 36 g/L](Mitchison-Field et al., 2019). For agar plates, 15g/L of agar was added to the media. Cells were grown in 5 mL of YPD with Instant Ocean broth at 22°C for 5 days unless otherwise specified.

**Table 1.** Place of origin of the *Hortaea* isolates.

| Strain | Species | Habitat |
| --- | --- | --- |
| CMF-20 | *Hortaea werneckii* | Coral, Woods Hole, MA |
| AMF-61 | *Hortaea werneckii* | Plankton Tow, Woods Hole, MA |
| EXF-120 | *Hortaea werneckii* | Hypersaline Water, Santa Pola saltpans, Spain |
| EXF-151 | *Hortaea werneckii* | Human, Tinea Nigra, Portugal |
| EXF-171 | *Hortaea werneckii* | Human; Keratomycosis (invasive); Brazil |
| EXF-562 | *Hortaea werneckii* | Soil on the sea coast, Namibia |
| EXF-2000 | *Hortaea werneckii* | Hypersaline Water, Secovlje saltpans, Slovenia |
| EXF-2682 | *Hortaea werneckii* | Human, Trichomycosis nigra (superficial); Italy |
| EXF-2788 | *Hortaea werneckii* | Hypersaline Water, Secovlje saltpans, Slovenia |
| EXF-4028 | *Salinomyces thailandica* | Glacier ice, Svalbard, Ny Alesund |
| EXF-6651 | *Hortaea werneckii* | Spider web, Atacama desert, Chile |
| EXF-6654 | *Hortaea werneckii* | Spider web, Atacama desert, Chile |
| EXF-6655 | *Hortaea werneckii* | Spider web, Atacama desert, Chile |
| EXF-6656 | *Hortaea werneckii* | Rock wall in a cave, Atacama desert, Chile |
| EXF-6669 | *Hortaea werneckii* | Spider web, Atacama desert, Chile |
| EXF-10513 | *Hortaea werneckii* | Deep See Water, Italy |

### Imaging EXF-2000

EXF-2000 cells were grown in 5 mL of Instant Ocean YPD broth at 30°C for 2-3 days. Cells were washed 3 times with 10 mL of PBS and resuspended with the corresponding YPD media. 100 μL of cell suspension (~1×10^6^ cells/mL) was inoculated in an Ibidi 15 μ-Slide I (Ibidi), and 1 mL of media was added to the reservoir. Cells were imaged for 18 hours, acquiring images every 5 minutes using a widefield microscope (Nikon Eclipse TI stage) with a Plan Apo λ 60x/1.40 Oil Ph3 DM objective and an Andor Zyla 4.2 plus VSC-06258 camera. The chi-square test was utilized to determine whether there was a statistically significant difference between the experimental groups using a *p-*value of 0.05 to determine significance. Since the overall test was significant, the maraschino procedure was used to determine where the difference occurred between the groups and avoid statistical significance attributable to performing multiple comparisons.

### *Macroscopic and Microscopic characterization of* Hortaea *isolates*

For macroscopic characterization, *Hortaea* isolates were streaked onto Instant Ocean YPD agar plates and incubated at 22°C for 5 days (Except EXF-4028 which was incubated for 10 days). For microscopic characterization, isolates were grown for 5 days in Instant Ocean YPD broth at 22°C. 20μl of cell solution was put onto a slide and imaged using a widefield microscope (Nikon Eclipse TI stage) with a Plan Apo λ 100x/1.40 Oil Ph3 DM objective and an Andor Zyla 4.2 plus VSC-06258 camera.

### Drug susceptibility testing

Isolates were grown in YPD+Instant Ocean agar for 7 days at 30°C. Cells were collected and washed 3 times with YPD. Cells were counted using a hemocytometer, and an initial stock solution of 1 x 10^7^ cells/ml of YPD was made. 3 1:10 serial dilution were performed to obtain the 1 x 10^6^, 1 x 10^5^, and 1 x 10^4^ cells/ml stocks. 10 μl of each isolate and their dilutions were plated into YPD, YPD+0.6M NaCl, YPD+1.7M NaCl, YPD+Fluconazole (4μg/ml), YPD+0.6M NaCl+Fluconazole (4μg/ml), YPD+Caspofungin (1μg/ml), and YPD+0.6M NaCl+Caspofungin (1μg/ml) agar. Plates were incubated at 30°C for 5 days.

### Galleria mellonella pathogenesis model and histology

Virulence of the *Hortaea* isolates was assessed by infecting 20 larvae of the wax moth *G. mellonella*. 5 μL of a 2×10^7^ cell / ml suspension for a total inoculum of 1×10^5^ cells per larvae or 5 μL of tissue-culture grade PBS were injected into the *G. mellonella* penultimate left proleg using a 10μl Hamilton syringe (catalog no. 80300) (Vargas-Muñiz et al., 2015). Infected larvae were incubated at 37°C and survival was scored every 24 hours for 5 days. Survival was plotted on a Kaplan-Meier curve with a log-rank pair-wise comparison. These experiments were repeated 3 times. Heat-killed EXF-2000 was done by incubating the yeast at 95°C for 1 hour. For histology, infected *G. mellonella* were harvested 1-day post-infection and injected with 100 μl of 10% buffered formalin (Beekman et al., 2018). Then, the larvae were fixed for 24 hours at 4°C in 5ml 10% buffered formalin. Larvae were cut into 3 transversal segments using a scalpel. These pieces were fixed for 48 hours at 4°C in 5ml 10% buffered formalin. Segments were embedded in paraffin and sectioned into 5 μm sections. Chromic Acid Schiff was performed on the sections and visualized via light microscopy.

## Results

### *H. werneckii* morphology and cell division pattern is modulated by NaCl concentration

In a previous study, only the condition that resembles the marine environment was evaluated. We tested if this method of growth was maintained in different conditions, or if *H. werneckii* alters its morphology and cell division pattern according to their environment (Mitchison-Field et al., 2019; Goshima, 2022). We used YPD, YPD supplemented with 0.6M NaCl (similar to ocean salinity), and YPD supplemented with 1.7M NaCl (similar to hypersaline salterns) and followed the growth of individual yeast cells for 18 hours at room temperature. When *H. werneckii* EXF-2000 isolate was grown in conditions that mimic marine salinity (YPD supplemented with 0.6M NaCl) we observed that EXF-2000 behaves similarly to the previously characterized *H. werneckii* isolate (Mitchison-Field et al., 2019). 75% of EXF-2000 grew as yeast, growing isotopically and from the poles before laying down septa (Figure 1A, 1C). Once a central septum is formed, EXF-2000 started budding from the poles almost simultaneously. Only 25% of the cells observed adopted a pseudohyphal-like morphology where multiple septa were formed, and few budding events were observed. 67% of cell division events observed were budding (Figure 1B). When EXF-2000 was grown on just YPD we observe that 65% of *H. werneckii* cells predominantly adopt a morphology that resembles that of pseudohyphae (Figure 1A, 1C). Concordant with the change in morphology, EXF-2000 predominantly divides via septation (Figure 1B). 64% of the cell division events observed were septation while budding only accounted for 36%. Growing EXF-2000 in YPD supplemented with 1.7M NaCl results in a morphology similar to the YPD+0.6M NaCl condition (Figure 1A-C).

**Figure 1.**
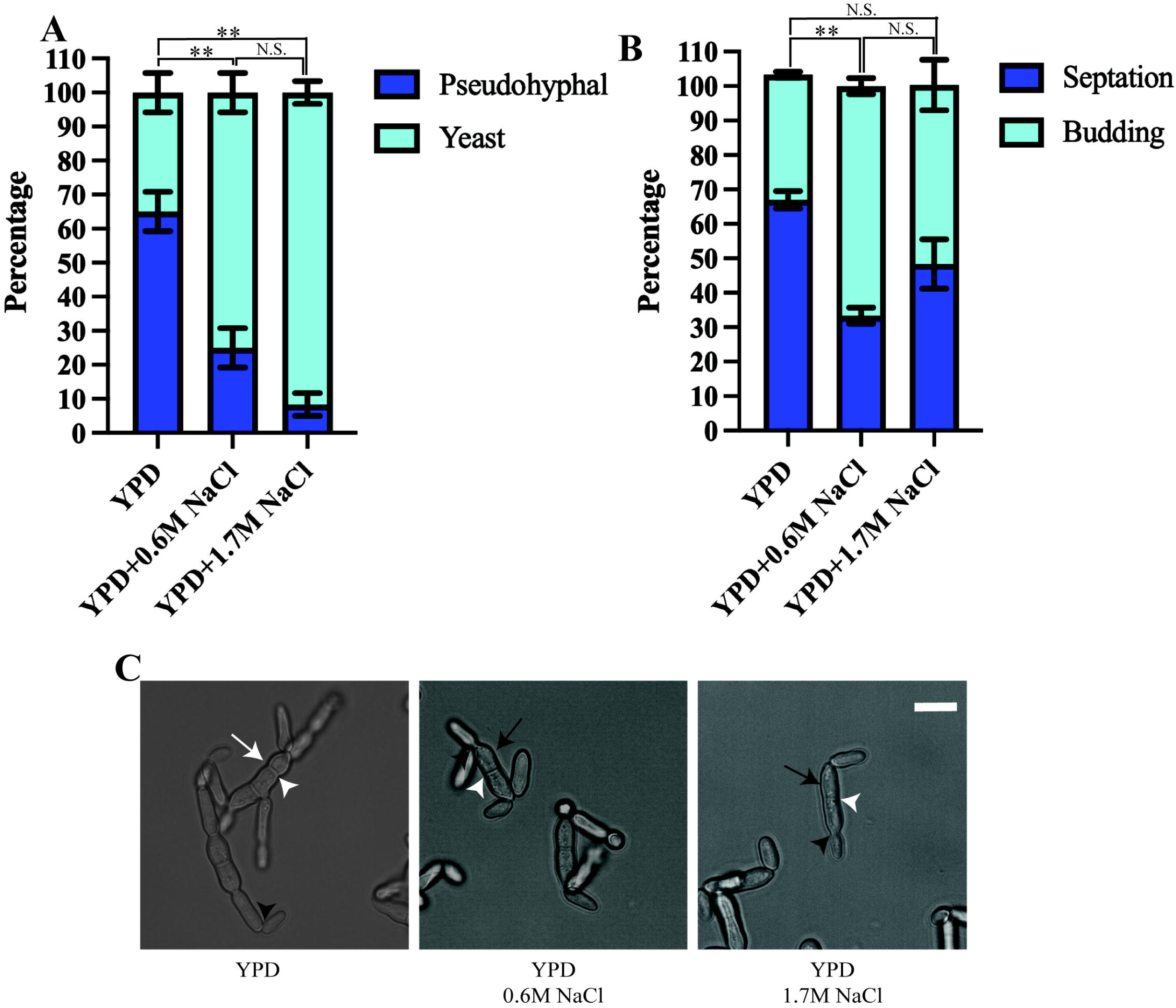
*Hortaea werneckii* EXF-2000 cell division and morphology is dependent on their osmotic environment. Experiments were conducted as 3 biological replicates, N=20 for each biological replicate. Error bars represent the standard error of the mean. Marascuilo procedure was used to determine statistical significance of individual conditions. Statistically significant comparison was represented by **. N.S. stands for not statistically significant. **(A)** EXF-2000 morphology distribution changes based on the presence of NaCl. (**B)** The preference of cell division mechanisms is altered when media is supplemented with NaCl. **(C)** Representative micrograph of EXF-2000 morphology after 18 hours of growth. White arrows point at pseudohyphal-like cells, while black arrows point to yeast cells. White arrowheads point at septa, while black arrowheads point at buds. Micrographs were obtained using a 60x objective. Scale bar,10μm

### *H. werneckii* isolates exhibit different colony and cellular morphology

Because *H. werneckii* has been isolated from different environments, we expect that different isolates will exhibit different morphotypes. We utilize previously isolated *H*. werneckii to test this hypothesis (Table 1). Most of the isolates have olive-green mucoid colonies by day 5 of growth in YPD supplemented with the Instant Ocean (Figure 2). EXF-171 and EXF-2682 exhibit a white-ish velvety growth indicative of hyphal growth. EXF-6656 also exhibit a velvet-like colony but the coloration was olive green. *Salinomyces thailandica* (EXF-4028) grew slower than the *H. werneckii* isolates. By day 5 only microcolonies of black coloration were observed (Data not shown). After 10 days of incubation, we start observing more sizable colonies of a rust color (Fig. 2).

**Figure 2.**
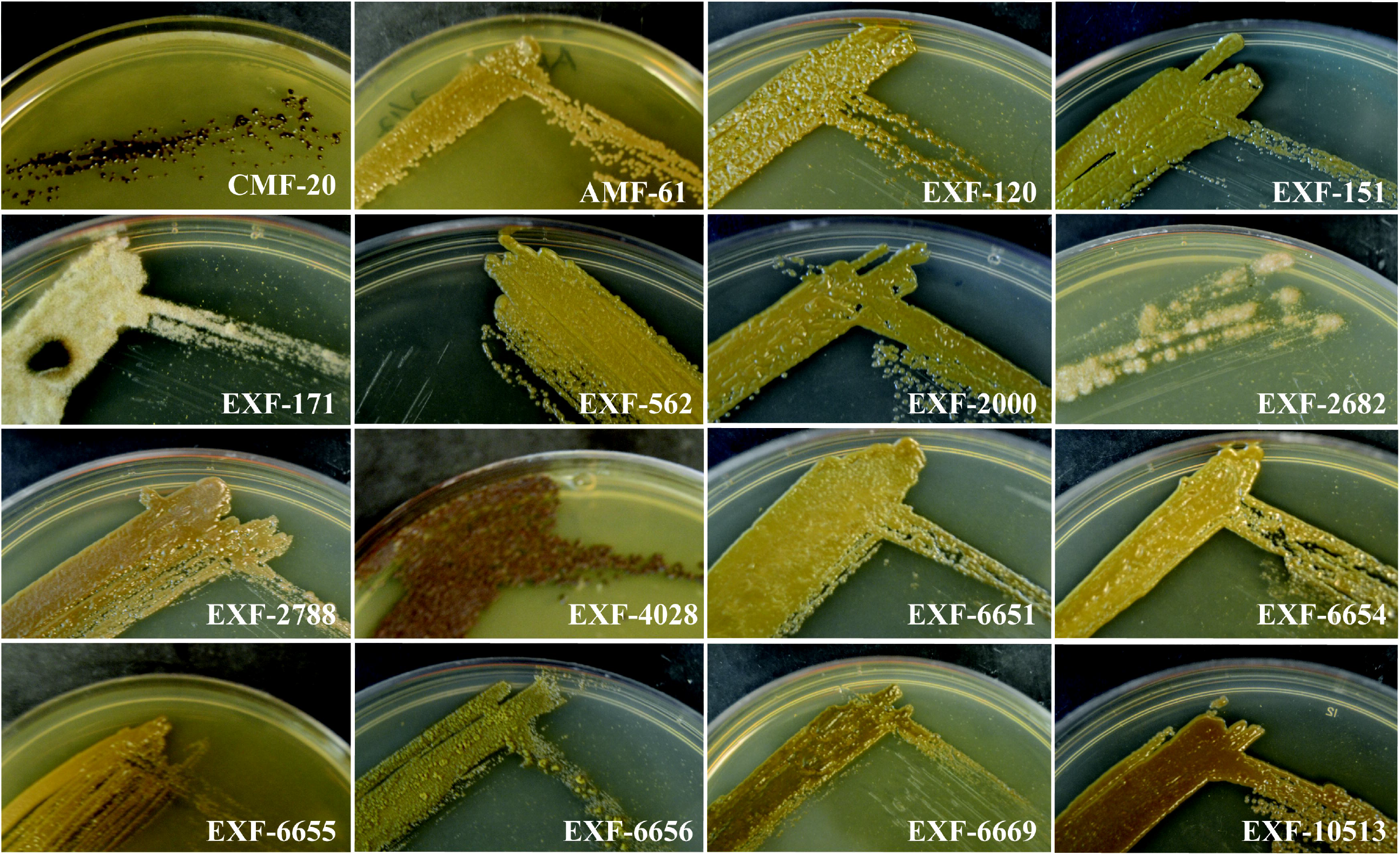
*Hortaea* isolates exhibit a great diversity in colony morphology and pigmentation. Isolates were grown for 5 days on Instant Ocean YPD, except for EXF-4028 that was grown for 10 days.

Similar to the colony morphology, the cell morphology was highly variable among the isolates (Figure 3). Most of the isolates grew as yeast cells in YPD broth supplemented with the Instant Ocean at 22°C. The yeast isolated from the Atacama Desert spider web seemed to be more round compared to other *H. werneckii* yeast. The CMF-20 isolate formed unique yeast cells that generate oblique and perpendicular septa. Only EXF-171, EXF-2682, and EXF-4028 formed true hyphae, with EXF-171 growing exclusively as hyphae.

**Figure 3.**
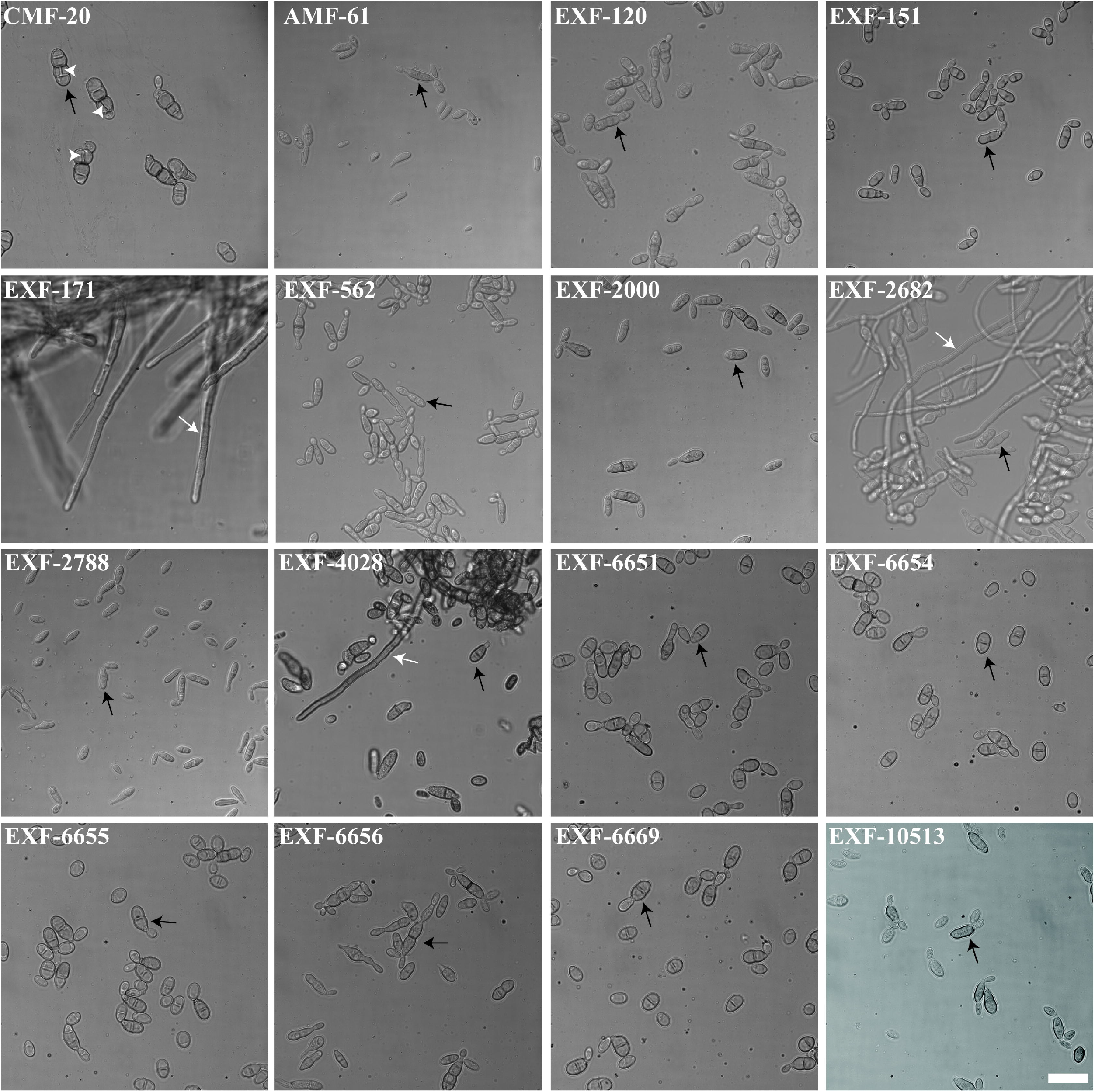
*Hortaea* isolates adopt different morphologies when grown in similar environmental conditions. Micrographs were obtained using a 100x objective. Scale bar, 20 μm. White arrows point at hyphae, while black arrows point to yeast cells, and white arrowheads point at transversal septation sites.

### *H. werneckii* isolates are differentially susceptible to fluconazole, but intrinsically resistant to caspofungin

Most *H. werneckii* isolates can grow in YPD+1.7M NaCl, with the notable exception of EXF-171. We observed that *H. werneckii* isolates grew better in YPD with fluconazole (4μg/ml) than in YPD+0.6M NaCl with fluconazole. Susceptibility to fluconazole was highly heterogeneous between each of the strains. Isolates EXF-120, EXF-2000, EXF-6655, and EXF-10513 showed higher susceptibility to fluconazole. EXF-6656 showed better resistance to fluconazole in the YPD+0.6M NaCl. Most *H. werneckii* were able to grow in the presence of caspofungin (1μg/ml). Only EXF-151, EXF-171, and EXF-562 showed susceptibility to the anti-cell wall drug. Nonetheless, all the isolates exhibited resistance to caspofungin when YPD was supplemented with caspofungin and 0.6M NaCl.

### *H. werneckii* isolates exhibit different pathogenic potential in an insect-based pathogenesis model

For many fungal pathogens, cell morphogenesis is linked to pathogenesis. Yeast cells usually correlate to dissemination, while hyphae are thought to facilitate invasion into host tissues in some fungal pathogens like *C. albicans* (Kornitzer, 2019). First, we tested if *H. werneckii* was capable of causing death in the *Galleria mellonella* insect model. To do this, we utilized different inoculum sizes of the EXF-2000 strain. We noted that using an inoculum size of 1 x10^5^ or 1 x 10^6^ cause 70% mortality or higher (Fig. S1) Since there is no statistical significance between the two inoculum sizes, we decided to utilize 1 x 10^5^ as our baseline to accommodate strains that grow poorly. Using this approach, we tested if *Hortaea* morphology correlates with pathogenesis. We infected 20 larvae with 1 x 10^5^ cells of each isolate and incubated them at 37°C. Only 6 out of the 16 isolates tested exhibited 40%mortality or higher after 5 days post-infection (Figure 5B), including the clinical isolates EXF-151 and EXF-2682. However, the EXF-171 clinical isolate only had a mortality rate of 40% or lower in our assays. The larvae’s death depended on fungal viability, as heat-killed yeast did not result in death (data not shown). Histopathology analyses showed that most of the pathogenic isolates had a yeast morphology in the host tissue (Figure 5C). The only exceptions were EXF-2682 and EXF-2000, which, in addition to yeast cells, had hyphae and pseudohyphae cells, respectively. Based on these results, we did not observe a direct correlation between hyphal morphology and virulence in our *Galleria* model.

**Figure 4.**
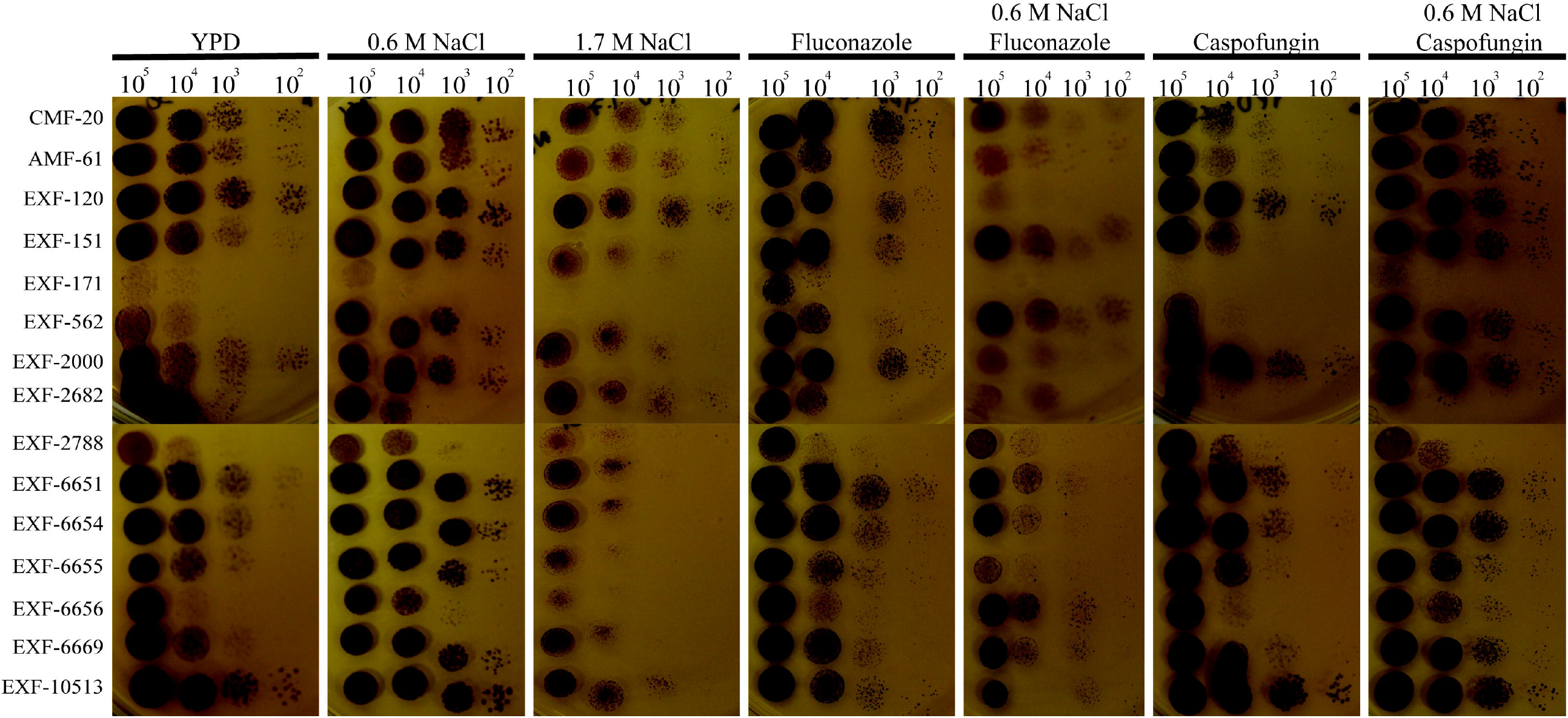
*H. werneckii* exhibits salt-dependent drug susceptibility. Different dilutions of *H. werneckii* cells were plated in YPD, YPD+0.6M NaCl, YPD+1.7M NaCl, YPD with fluconazole (4μg/ml), YPD+0.6M NaCl with fluconazole, YPD with caspofungin (1μg/ml) and YPD+0.6M NaCl with caspofungin. Plates were incubated at 30°C for 5 days. *H. werneckii* were more susceptible to fluconazole when grown in YPD+0.6M NaCl, while they were more susceptible to caspofungin when grown only in YPD.

**Figure 5.**
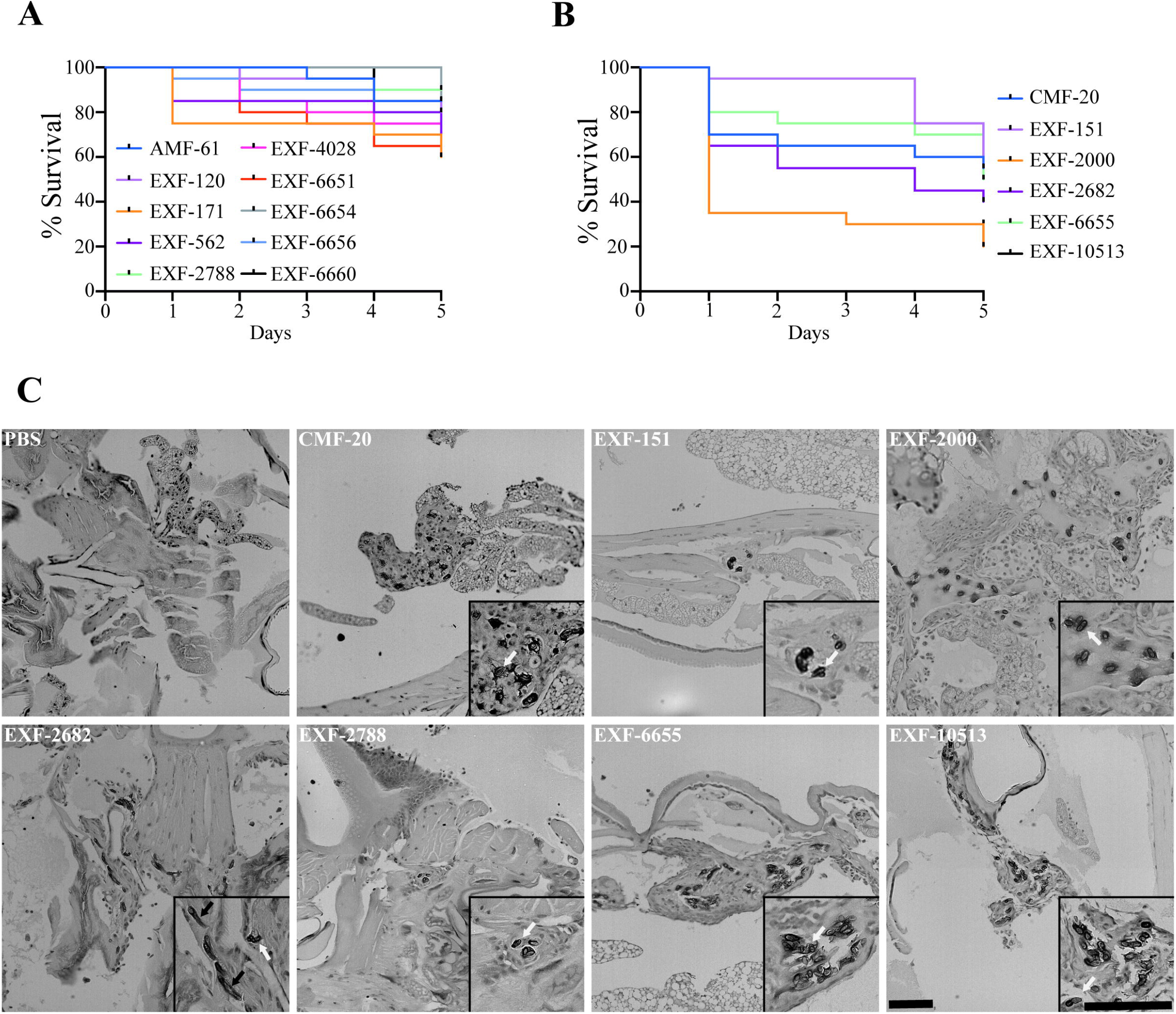
Only a subset of the isolates can cause more than 40% mortality in the *Galleria mellonella* insect model. (A) Isolates that caused <40% mortality after 5 days post-inoculation (DPI). (B) Isolates that caused >40% mortality after 5 DPI. (C) Histopathology of infected *G. mellonella* after 1 day post-infection. Scale bar, 100 μm. 20X

## Discussion

*Hortaea werneckii* is a black yeast commonly found in extreme environments, such as salterns, the ocean, and the desert. Previous studies revealed that *H. werneckii* has a unique cell division cycle (Mitchison-Field et al., 2019). It begins as a yeast cell that grows in a manner that resembles that of the fission yeast. After it lays down septa, *H. werneckii* starts budding from its pole in a fashion that is like that of budding yeast. These cell division programs and morphologies are dependent on *H. werneckii* cell density (Goshima, 2022). Here, we demonstrated that this growth pattern is dependent on the environment in which *H. werneckii* is growing. This is in agreement with previous microarray data that shows that *H. werneckii* regulates the expression of cell cycle and cell wall-related genes in response to the hypersaline environment (Petrovič et al., 2002; Vaupotix and Plemenitaš, 2007). A subset of these genes was dependent on the Hog1 osmolarity response pathway. In *C. albicans*, Hog1 inhibits the yeast-to-hyphae transition (Eisman et al., 2006). In *H. werneckii* Hog1 is activated at high osmolarity, possibly contributing to the yeast morphology. When *H. werneckii* cells are growing in YPD, it is possible that the Hog1 pathway is inactive leading to the yeast-to-pseudohyphal transition we observe in the EXF-2000 strain (Kejžar et al., 2015). However, previous studies using a Hog1 inhibitor only observed the yeast morphology. This points to Hog1 possibly being insufficient for the changes in morphology observed in *H. werneckii* EXF-2000.

*Hortaea spp*. have been isolated from a great range of environments varying in salinity. To understand if the environment affected the morphology observed, we characterize the colony morphology as well as their cellular morphology. All of the strains were capable of growing in conditions similar to the ocean. This indicates that regardless of where the *Hortaea* strains were isolated, these isolates can still adapt to high salinity. Two of the three clinical isolates exhibited true hyphal growth under the experimental condition we utilized. Hyphae are critical for tissue invasion in many fungal pathogens, and it might be a morphology that is selected for in pathogenic *H. werneckii*. However, more clinical isolates need to be analyzed to determine if the ability to form hyphae is important in *H. werneckii* infection. The majority of the candidates analyzed grew as yeast with a central septum as previously described(Mitchison-Field et al., 2019; Zalar et al., 2019). However, some of the CMF-20 cells would form a septum that is perpendicular to the previous septation sites. This septation pattern is reminiscent of the septation pattern of *Neophaeotheca salicorniae*, a *Capnodiales* yeast associated with saline environments (Mitchison-Field et al., 2019). However, to our knowledge, no other *H. werneckii* isolate exhibits this septation pattern.

None of the isolates were susceptible to the anti-cell wall drug caspofungin when grown in YPD+0.6M NaCl and 1μg/ml of caspofungin. Similar to our findings, previous studies have shown that *H. werneckii* displays various levels of susceptibility to caspofungin when the media is not supplemented with salt (Badali et al., 2019). In the presence of salt, *H. werneckii* cells upregulated pumps and utilize melanin and osmolytes to adapt to osmotic stress. High melanization of the cells is observed when cells are grown in the presence of caspofungin. Melanin helps with the retention of glycerol in *H. werneckii* (Kogej et al., 2007). This robust osmoadaptation mechanism may be behind the observed *H. werneckii* resistance to cell wall stress caused by caspofungin.

Contrary to caspofungin, *H. werneckii* was more susceptible to fluconazole when grown in YPD+0.6M NaCl. Fluconazole inhibits ergosterol synthesis, increasing the plasma membrane fluidity and permeability (Abe et al., 2009). In *S. cerevisiae*, salt stress results in higher ergosterol content (Turk et al., 2004). Similarly, it is possible that at higher salt concentrations, *H. werneckii* relies more heavily on ergosterol to maintain the physical properties of its plasma membrane. This would result in higher susceptibility to fluconazole. However, previous studies contradict this notion (Turk et al., 2004). Contrary to *S. cerevisiae*, the ergosterol content of *H. werneckii* is not significantly altered as salinity is increased. Similarly, membrane fluidity is not altered by an increase in salinity.

*H. werneckii* is the etiological agent of tinea nigra. Tinea nigra manifests as dark spots in the sole of feet or the palm of the hand of immunosuppressed individuals (Bonifaz et al., 2008; Giordano Lorca et al., 2018). Due to the morphological diversity observed in the *Hortaea* isolates, we decided to test if morphology correlated with pathogenic potential of this fungal pathogen using a *Galleria mellonella* insect model. Using this model, we observed that only 6 of these isolates were able to cause >40% mortality rates in our experimental conditions, including two clinical isolates (EXF-2682 and EXF-151). Nonetheless, morphology by itself did not predict higher virulence of the isolates, indicating that there are other factors that contribute to *Hortaea* pathogenesis. Based on the previous reports, some of the factors that could be key for pathogenicity is the ability to grow at 37°C and the production of proteases, as the strains with higher virulence exhibiter these adaptations (Zalar et al., 2019). Additionally, there is no phylogenetic relationship between the isolates with high virulence indicating that the ability to establish infection might have evolved multiple times in the *Hortaea* lineage (Gostinčar et al., 2018).

The *G. mellonella* system can help understand the pathogenic potential of different isolates, providing a better understanding of tinea nigra and disseminated phaeohyphomycosis. These analyses can be coupled with genomics studies to identify the mechanism of host colonization and virulence that could lead to improved patient outcomes. Nonetheless, the *G. mellonella* system fails to fully recapitulate the clinical presentations of severe disseminated phaeohyphomycosis. For example, this system relies on immunocompetent larvae; however, patients with severe phaeohyphomycosis have underlying immune disorders such as *CARD9* mutations (Bonifaz et al., 2013; Wang et al., 2014, 2020; Vaezi et al., 2018). As more tools and isolates of *H. werneckii* become available, developing a murine model of disseminated phaeohyphomycosis will be necessary to test the possible clinical intervention in a system with similar biology to the patients.

*Hortaea werneckii* is of particular interest due to most isolates originated from intraspecific hybridization (Gostinčar et al., 2018; Zalar et al., 2019; Romeo et al., 2020). For example, EXF-2000 and EXF-120 lineages originated from an intraspecific hybridization of the EXF-562 and EXF-2788 lineage (Gostinčar et al., 2018). Nonetheless, only the EXF-2000 is more virulent in our *Galleria* model, which could be a phenotype that arose from intraspecific hybridization (Zalar et al., 2019). Fungi, including human pathogens, are capable of hybridization, generating significant genomic and phenotypic diversity (Mixão and Gabaldón, 2018, 2020; Steenwyk et al., 2020; Theelen et al., 2022). This is most likely the case with *H. werneckii*, where these intraspecific hybridization events provide *Hortaea* species with the genomic tools needed to adapt to extreme environments and the human host. This is further evident as the 6 isolates with the higher pathogenic potential are all intraspecific hybrids (Gostinčar et al., 2018, 2022).Characterization of more isolates coupled with genomic analyses will further our understanding of the contribution of intraspecific hybridization to the phenotypic plasticity observed in *H. werneckii* and the evolution of fungal pathogenic traits.

## Acknowledgments

The authors want to thank Dr. Nina Gunde-Cimerman and Dr. Amy Gladfelter for sharing their isolate collection with us. The authors want to also thank Dr. Connie Nichols (Duke) and Dr. Andrew Alspaugh (Duke) for technical advice. Histological services were provided by the Histology Research Core Facility in the Department of Cell Biology and Physiology at the University of North Carolina at Chapel Hill. JVM wants to thank the Gladfelter lab (UNC), William Wolf (UNC), Fisher Lab (SIU) and the Vargas-Muñiz Lab for critical reading of this manuscript.

---

**Figure S1. Establishing an insect-based model to assess *H. werneckii* virulence**. Different concentration of the EXF-2000 strain were injected into 10 *Galleria mellonella* larvae. Larvae were incubated at 37°C and survival was monitored for 5 days.

## Notes

### Competing Interest Statement

The authors have declared no competing interest.

