## Supplementary figures and images for "*Hortaea werneckii* isolates exhibit different pathogenic potential in the invertebrate infection model *Galleria mellonella*"

### Supplemental Figure 1

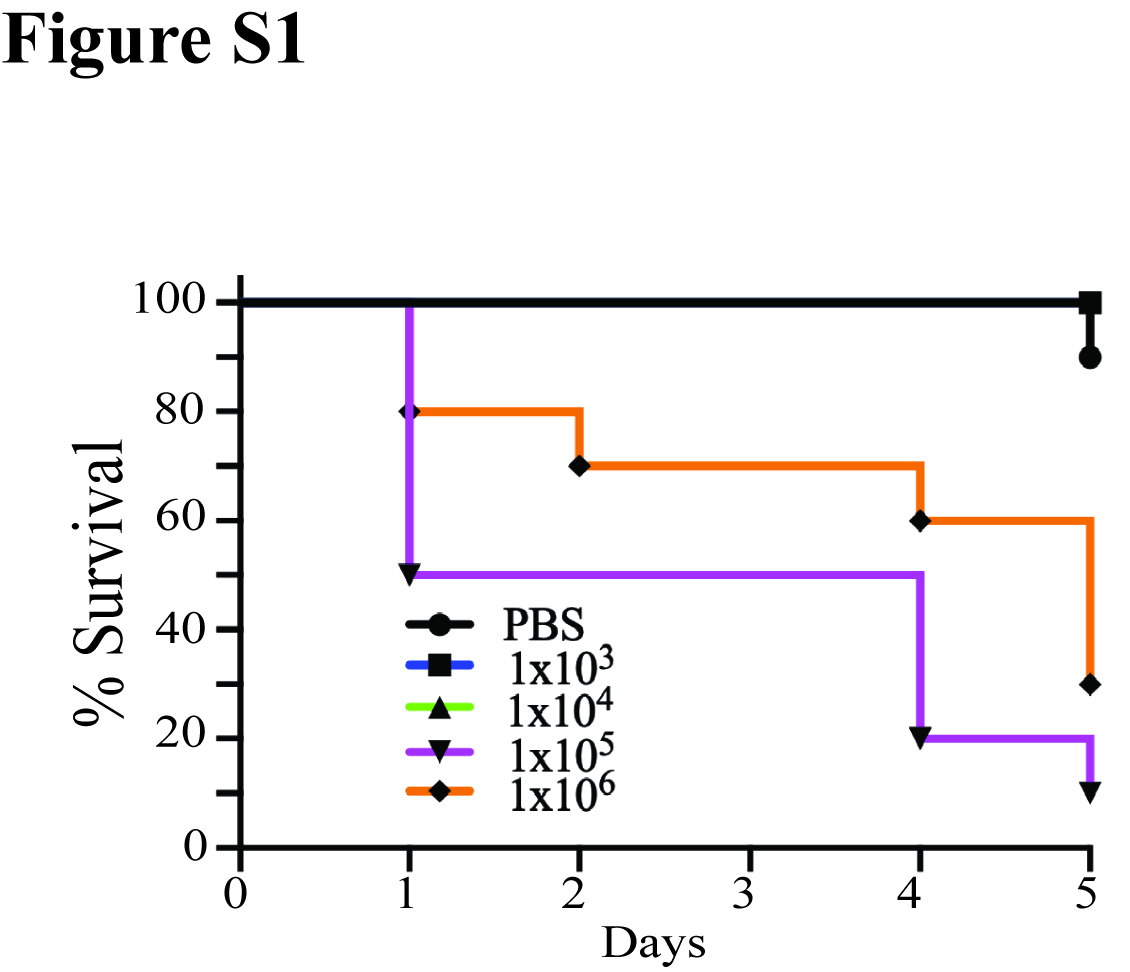
